# Block of Thalamic 5-HT2A Receptors modulates Tonic GABA-A Inhibition but not Absence Seizures

**DOI:** 10.64898/2026.09.19.751734

**Authors:** Anna Cavaccini, Marcello Venzi, Vincenzo Crunelli, Giuseppe Di Giovanni

## Abstract

Absence epilepsy is a generalized non-convulsive epilepsy characterized by recurrent absence seizures (ASs) and enhanced GABAA receptor-mediated tonic inhibition of thalamocortical (TC) neurons has been shown to lead to ASs. Serotonergic signalling is an important modulator of these seizures, yet the contribution of specific 5-HT receptor subtypes remains incompletely understood. Here, we investigated whether endogenous serotonergic signalling through 5-HT2A receptors (5-HT2ARs) contributes to pathological tonic inhibition and ASs expression. Patch-clamp recordings were obtained from ventrobasal (VB) TC neurons of Genetic Absence Epilepsy Rats from Strasbourg (GAERS) and non-epileptic control (NEC) rats. The selective 5-HT2AR antagonist MDL11,939 had no effect on tonic GABA-A currents in NEC TC neurons but significantly reduced the enhanced tonic current in GAERS neurons, revealing a disease-dependent contribution of endogenous 5-HT2AR signalling. MDL11,939 administered bilaterally into the VB of freely moving GAERS by reverse microdialysis did not alter ASs. This dissociation between the modulation of a disease-associated cellular mechanism and seizure outcome suggests that the previously reported reduction of ASs by systemic 5-HT2AR blockade may involve mechanisms outside the VB.

---

Typical absence seizures (ASs) arise from abnormal interactions between the cerebral cortex and thalamus, resulting in highly synchronized spike-and-wave discharges (SWDs) associated with transient impairment of consciousness (Crunelli and Leresche, 2002; Crunelli et al., 2020). A hallmark of ASs is enhanced GABA-A receptor-mediated tonic inhibition in thalamocortical (TC) neurons of the ventrobasal (VB) thalamus. This tonic conductance is markedly increased in VB TC neurons of Genetic Absence Epilepsy Rats from Strasbourg (GAERS) due to impaired astrocytic GABA transporter-1 (GAT-1)-mediated uptake and contributes to the pathological thalamocortical synchronization underlying SWDs (Cope et al., 2009).

Serotonin (5-HT) is an important modulator of thalamocortical excitability, and several 5-HT receptor subtypes have been implicated in AS regulation. In GAERS, activation of 5-HT2C receptors (5-HT2CRs) with Ro 60-0175 reduces tonic GABA-A currents in VB TC neurons and markedly decreases spontaneous ASs, providing a potential cellular mechanism for its anti-absence action (Cavaccini et al., 2026).

The actions of 5-HT2A receptors (5-HT2ARs) appear more complex and depend on the animal model, receptor localization, and network state. 5-HT2ARs can regulate thalamic inhibitory transmission through TC and reticular thalamic neurons (Goitia et al., 2016) and astrocytes (Crunelli et al., 2026). In normal Wistar rats, activation of 5-HT2ARs with TCB-2 reduces GABA uptake, enhances tonic GABA-A currents in VB TC neurons, and induces ethosuximide-sensitive ASs following bilateral administration into the VB, an effect primarily mediated by astrocytic 5-HT2AR-dependent inhibition of GAT-1-mediated GABA uptake (Crunelli et al., 2026). In contrast, in GAERS, TCB-2 reduces AS expression, whereas systemic administration of the selective 5-HT2AR antagonist MDL11,939 produces only a transient increase in the total time spent in seizures (Venzi et al., 2016). Moreover, local infusion of TCB-2 into the VB suppresses ASs (Morais et al., 2026), and evidence indicates that 5-HT2AR activation in VB TC neurons reduces tonic GABA-A inhibition (Deidda et al., 2019). These apparently contrasting findings raise the question of how endogenous 5-HT signalling through 5-HT2ARs contributes to pathological tonic inhibition and AS expression in GAERS.

We therefore investigated whether endogenous 5-HT2AR activity contributes to tonic GABA-A inhibition in VB TC neurons from NEC and GAERS rats and whether local blockade of these receptors affects spontaneous Ass (see Supplementary Methods). Whole-cell voltage-clamp recordings were obtained from visually identified VB TC neurons in thalamic slices from 25–30-day-old male NEC and GAERS rats. Tonic GABAA current was quantified as the gabazine-sensitive shift in holding current normalized to whole-cell capacitance. Endogenous 5-HT2AR signalling was assessed using MDL11,939 (500 nM). For in vivo experiments, freely moving 3–4-month-old male GAERS were implanted with cortical EEG electrodes and bilateral microdialysis probes targeting the VB. Vehicle or MDL11,939 (50 μM) was administered by reverse microdialysis, and seizure number, mean duration, and cumulative seizure time were quantified.

Under control conditions, the tonic GABAA current was larger in TC neurons from GAERS than NEC rats (Fig. 1A1-A3), confirming the pathological enhancement of tonic inhibition in this rat model (Cope et al., 2009). MDL11,939 did not significantly alter tonic current in NEC neurons, consistent with observations in Wistar rats (Crunelli et al., 2026), indicating that endogenous 5-HT2AR activity does not detectably contribute to physiological tonic inhibition under these *ex vivo* conditions. Unexpectedly, MDL11,939 reduced tonic GABAA current in GAERS TC neurons (Fig. 1B1-B3), indicating that endogenous 5-HT2AR signalling contributes to the enhanced tonic inhibition associated with the epileptic phenotype. Despite this cellular effect, bilateral administration of MDL11,939 into the VB did not alter the cumulative seizure time, seizure number and mean duration (Fig. 1C2–C4).

**Figure 1.**
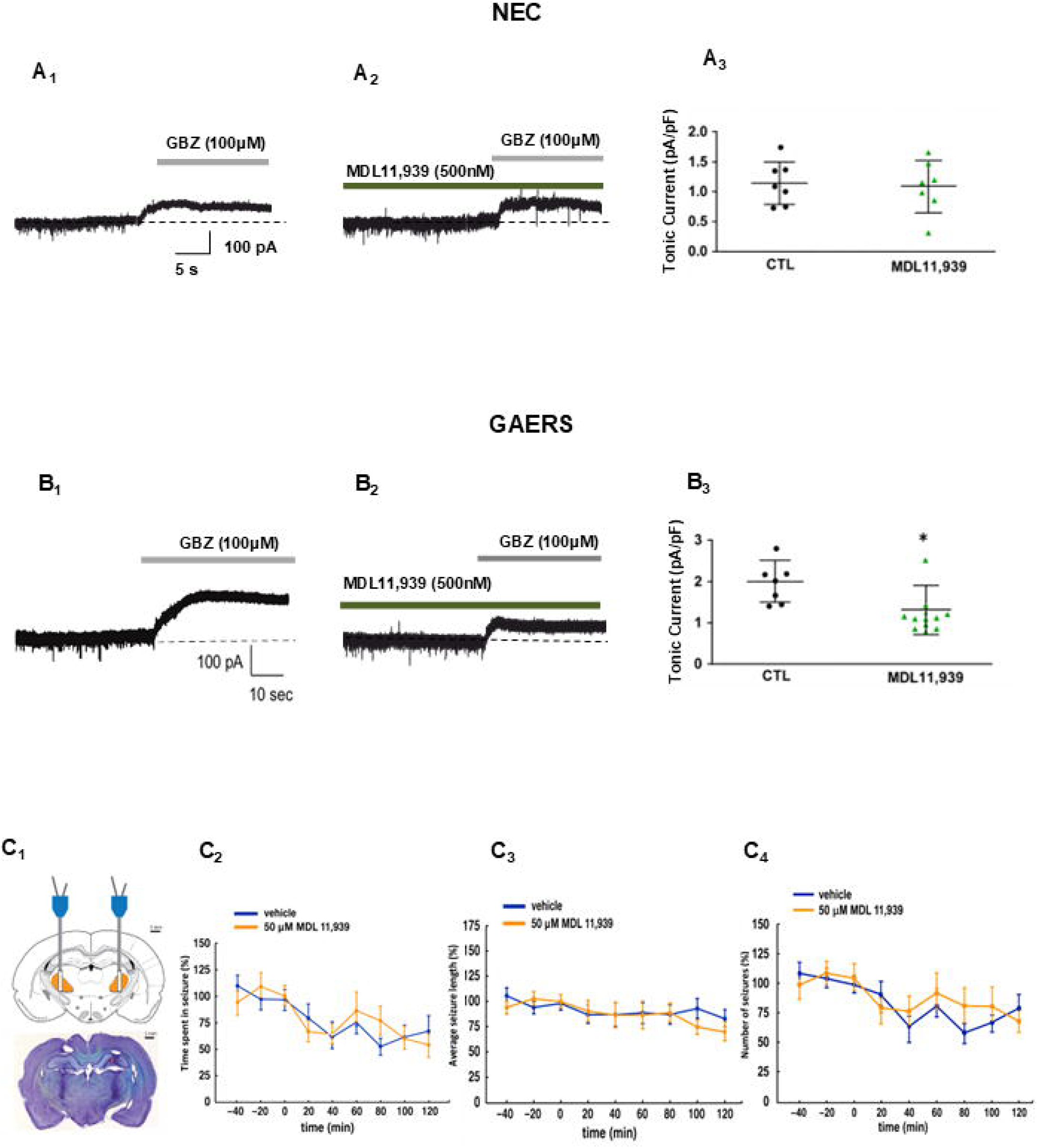
Selective of thalamic 5-HT2ARs reduces tonic GABA-A currents in GAERS *in vitro* but does not affect their ASs *in vivo*. **(A1-A3)** Representative whole-cell voltage-clamp recordings from VB TC neurons of non-epileptic control (NEC) rats showing the gabazine (GBZ, 100 μM)-sensitive tonic GABA-A current under control conditions (**A1**) and in the presence of the selective 5-HT2AR antagonist MDL11,939 (500 nM) (**A2**). **A3**, Summary scatter plot of tonic current (normalized to cell capacitance) showing that MDL11,939 does not affect tonic GABA-A current in NEC TC neurons. Control group data were obtained from Cavaccini et al. (2026), as the experiments involving MDL11,939 were performed in the same experimental cohort. **(B1–B3)** Representative whole-cell voltage-clamp recordings from VB TC neurons of GAERS under control conditions (**B1**) and during bath application of MDL11,939 (500 nM) (**B2**). **B3**, Summary scatter plot showing that MDL11,939 significantly reduced tonic GABA_A_ current in GAERS TC neurons. In A3 and B3, individual symbols represent single neurons, and horizontal bars indicate mean ± SEM. *P* < 0.05 versus control (CTL). Control group data were obtained from Cavaccini et al. (2026), as the experiments involving MDL11,939 were performed in the same experimental cohort. **(C1–C4) C1**, Top panel: schematic illustration of bilateral microinjection sites within the VB thalamus. Bottom panel: representative coronal section confirming injection placement. **C2–C4**, Time course of ASs parameters before and during bilateral intra-thalamic administration of vehicle or MDL11,939 (50 μM). **C2**, total time spent in seizures; **C3**, average seizure duration; and **C4**, number of seizures. Drug administration is from time zero onwards. Data are expressed as mean ± SEM. Intra-thalamic administration of MDL11,939 did not significantly alter any seizure parameter compared with vehicle-treated animals.

Thus, the transient pro-absence effect previously observed following systemic MDL11,939 administration (Venzi et al., 2016) is unlikely to result from 5-HT2AR blockade within the VB and cannot be attributed to an increase in tonic GABAA inhibition in VB TC neurons. Intriguingly, both blockade of endogenous 5-HT2AR signalling in the present study and receptor activation with TCB-2 (Deidda et al., 2019) reduce tonic GABA-A inhibition in GAERS. This apparent paradox suggests that a simple bidirectional relationship between 5-HT2AR activity and tonic GABAA inhibition cannot explain these findings. Instead, the effects may depend on the recruitment of distinct 5-HT2AR populations on TC neurons, astrocytes, and reticular thalamic terminals, as well as differences between endogenous receptor tone and pharmacological activation.

The lack of effect of VB-infused MDL11,939 on ASs may also reflect the more limited impact of removing endogenous 5-HT2AR tone compared with receptor activation and/or limited receptor engagement and tissue coverage achieved by reverse microdialysis. Alternatively, the tendency toward a pro-absence effect of systemic 5-HT2AR blockade may reflect actions outside the VB, particularly in the somatosensory cortex, a key region for SWD initiation that expresses high levels of 5-HT2ARs (Crunelli et al., 2020). Cortical 5-HT2AR blockade could therefore facilitate mechanisms involved in AS initiation, although this hypothesis requires direct experimental verification.

Overall, these results reveal a state- and circuit-dependent role of 5-HT2ARs in absence epilepsy and a dissociation between their contribution to pathological thalamic tonic GABA-A inhibition and the control of AS expression. The pro-absence effect of systemic MDL11,939 may therefore involve extra-thalamic mechanisms, potentially within cortical networks.

## CRediT authorship contribution statement

**Anna Cavaccini:** Investigation; methodology; data analysis; figure preparation; original draft. **Marcello Venzi**: Investigation; methodology; data analysis; figure preparation; original draft. **Vincenzo Crunelli:** Conceptualization; study design; supervision; funding acquisition; manuscript drafting and editing. **Giuseppe Di Giovanni:** Conceptualization; study design; supervision; funding acquisition; data analysis; manuscript drafting and editing.

## Acknowledgements

We wish to thank Mr. Timothy Gould for technical assistance.

## Data availability

Data are available from the corresponding author upon reasonable request.

