## Supplementay Materials & Methods for "Block of Thalamic 5-HT2A Receptors modulates Tonic GABA-A Inhibition but not Absence Seizures"

**Supplementary MATERIALS AND METHODS**

**Animals and ethical statement**

Male Genetic Absence Epilepsy Rats from Strasbourg (GAERS) and non-epileptic control (NEC) rats were bred in-house at the School of Biosciences, Cardiff University (UK). Animals were housed under controlled environmental conditions (19–21 °C; 45–65% relative humidity) with food and water available *ad libitum*. Postnatal day (P) 25–30 rats were used for the *in vitro* electrophysiological experiments, whereas adult male GAERS (3–4 months old) were used for the *in vivo* EEG and reverse microdialysis experiments, consistent with the experimental populations described in the study.

All experimental procedures were conducted at Cardiff University under UK Home Office and Cardiff University ethical approval and in accordance with the UK Animals (Scientific Procedures) Act 1986 and institutional guidelines for animal welfare. All efforts were made to minimize animal suffering and the number of animals used.

***In vitro* electrophysiology**

Horizontal brain slices containing the ventrobasal (VB) thalamus and nucleus reticularis thalami (NRT) were prepared as previously described (Cope et al., 2009). Rats were deeply anesthetized with isoflurane and the brain was rapidly removed. Slices (300 μm thick) were prepared in ice-cold solution continuously oxygenated with 95% O₂/5% CO₂ and containing (in mM): 60 sucrose, 85 NaCl, 25 NaHCO₃, 2.5 KCl, 1.25 NaH₂PO₄, 1 CaCl₂, 2 MgCl₂, 10 D-glucose and 3 kynurenic acid (approximately 300 mOsm, pH 7.4).

Slices were incubated for 20 min at 32 ± 0.5 °C in the same solution without kynurenic acid and then transferred for a further 40 min to sucrose-free artificial cerebrospinal fluid (aCSF) containing (in mM): 125 NaCl, 25 NaHCO₃, 2.5 KCl, 1.25 NaH₂PO₄, 1 CaCl₂, 2 MgCl₂ and 10 D-glucose.

Following recovery, individual slices were transferred to the recording chamber and continuously perfused at approximately 2 ml/min with oxygenated aCSF maintained at 32 ± 0.5 °C. Recording aCSF contained 2 mM CaCl₂, 1 mM MgCl₂, 0.5 μM tetrodotoxin (TTX) and 3 mM kynurenic acid. Experiments were performed on only one neuron per slice.

**Whole-cell patch-clamp recordings**

Thalamocortical (TC) neurons of the VB were visually identified using a Nikon Eclipse E600FN microscope (Nikon, UK) equipped with a 40× water-immersion objective and video camera (Hamamatsu, UK). Whole-cell voltage-clamp recordings were obtained from TC neurons held at −70 mV using glass patch pipettes with a resistance of 2.5–3.5 MΩ. Pipettes contained an intracellular solution composed of (in mM): 130 CsCl, 2 MgCl₂, 4 Mg-ATP, 0.3 Na-GTP, 10 Na-HEPES and 0.1 EGTA (pH 7.35; approximately 305 mOsm).

Series resistance and whole-cell capacitance were determined from responses to 5-mV voltage steps. Series resistance was compensated by 70–80% and continuously monitored throughout the recordings. Recordings were excluded if access resistance, initially 4–10 MΩ, changed by more than 20%.

Tonic GABAA receptor-mediated current was measured as the shift in holding current produced by application of the GABA${}_{A}$ receptor antagonist gabazine (GBZ, 100 μM), as previously described (Cope et al., 2009). GBZ was applied after a stable baseline had been established. Baseline current was calculated as the average current during the 10 s immediately preceding GBZ application. The GBZ-sensitive shift in holding current was determined from the average current during a 10-s period following GBZ application or at the peak of the stable baseline shift. Tonic GABAA current amplitude was normalized to whole-cell capacitance and expressed as pA/pF.

To investigate the contribution of endogenous 5-HT₂A receptor signalling, slices were perfused with the selective 5-HT₂A receptor antagonist MDL11,939 (500 nM) for at least 10 min before GBZ application. Corresponding vehicle conditions were used for control recordings. The manuscript reports MDL11,939 at 500 nM for these experiments.

**Surgical implantation of EEG electrodes and microdialysis guide cannulae**

Adult male GAERS were anesthetized with isoflurane. General anesthesia was induced with 5% isoflurane in 2 L/min O₂. Animals were then transferred to a stereotaxic frame and anesthesia was maintained with isoflurane in O₂, initially at approximately 3.5% and progressively reduced to approximately 2% during surgery. Adequate anesthesia was confirmed by the absence of hind-limb withdrawal and tail-pinch reflexes. Body temperature was monitored using a rectal probe and maintained at approximately 37 °C with a homeothermic heating system.

Sterile saline (0.9%, 5 ml/kg, s.c.) was administered to maintain hydration, and meloxicam (1 mg/kg, s.c.) was administered for postoperative analgesia. Animals were monitored during recovery and received additional postoperative care when required. At least 5 days of recovery were allowed before experimental recordings.

For chronic EEG recording, holes were drilled bilaterally over the frontal and parietal cortices for placement of epidural EEG electrodes. Additional screws were used to stabilize the implant. EEG electrodes consisted of gold-plated screws connected to insulated copper wires and a head-mounted connector. The electrodes and anchor screws were secured with dental cement while leaving the *dura mater* intact.

For reverse microdialysis experiments, two additional holes were drilled for bilateral implantation of guide cannulae targeting the VB thalamus. Stereotaxic coordinates were AP −3.4 mm and ML ±2.8 mm relative to bregma. Guide cannulae for CMA 12 microdialysis probes were slowly lowered into position, with a final DV position of −4.4 mm measured from the bottom of the guide cannula. The guide cannulae were secured with acrylic cement and fitted with dummy probes until the experimental session.

**Freely moving EEG recordings and reverse microdialysis**

For reverse microdialysis experiments, dummy probes were removed and two CMA 12 MD Elite microdialysis probes with a 2-mm membrane were slowly inserted into the guide cannulae. Experiments were performed 18–24 h after probe insertion. Rats were placed individually in recording chambers and connected to the EEG recording system while remaining freely moving.

Microdialysis probes were connected through FEP tubing to syringes mounted on a CMA 400 syringe pump. Perfusion was maintained at 1 μl/min. Before drug administration, animals were allowed to habituate to the recording chamber while the probes were continuously perfused with vehicle.

Following the baseline period, the perfusion solution was either maintained as vehicle or switched to MDL11,939 (50 μM), which was administered bilaterally into the VB by reverse microdialysis. EEG activity was continuously recorded before and during treatment. The present study quantified seizure activity during bilateral intra-VB administration of vehicle or MDL11,939.

At the end of each recording session, the microdialysis probes were removed and replaced with clean dummy probes. A minimum interval of 6 days was allowed between consecutive recording sessions.

**Histological verification of microdialysis probe placement**

At the completion of the experiments, the brains were collected for histological verification of microdialysis probe placement. Coronal brain sections were examined to confirm that the dialysis probes had been positioned within the VB thalamus. Experiments in which probe placement was outside the intended target region were excluded from the analysis. The location of the bilateral VB administration sites and representative histological verification are shown in Figure 1C1 of the manuscript.

**Detection of spike-and-wave discharges**

Spike-and-wave discharges (SWDs) were detected from cortical EEG recordings using the SeizureDetect procedure and subsequently verified by visual inspection. Raw EEG signals were DC-corrected, and a baseline epoch of desynchronized waking EEG was used as the control period.

Candidate SWDs were initially identified using an amplitude threshold defined relative to the mean and standard deviation of the control EEG. Temporal criteria were subsequently applied to identify the onset and continuation of individual SWDs, including the interval between successive peaks, minimum event duration and minimum interval between separate events. Candidate events were further classified according to their characteristic oscillatory frequency, thereby allowing SWDs to be distinguished from sleep-related oscillations and recording artefacts. Automated detection was subsequently refined by visual inspection.

**Quantification of absence seizures**

Absence seizures were quantified using three parameters: cumulative time spent in seizures, mean seizure duration and total number of seizures. These correspond to the three outcome measures used to assess the effect of intra-VB MDL11,939 in the present study.

Seizure parameters were quantified in consecutive 20-min epochs. Values obtained during the treatment period were normalized to the corresponding pre-treatment control period to reduce inter-animal variability and allow comparison of the time course of drug effects. The resulting normalized values were used to compare vehicle and MDL11,939 treatment conditions.

**Drugs and solutions**

MDL11,939 was used as the selective 5-HT₂A receptor antagonist throughout the study. For *in vitro* experiments, MDL11,939 was applied at a final concentration of 500 nM. For *in vivo* experiments, MDL11,939 was administered bilaterally into the VB by reverse microdialysis at 50 μM.

Gabazine (GBZ, 100 μM) was used to determine the GABA${}_{A}$ receptor-mediated tonic current. Tetrodotoxin (TTX, 0.5 μM) and kynurenic acid (3 mM) were included in the recording aCSF to isolate inhibitory currents. MDL11,939 was dissolved in ethanol and diluted to the required final concentration immediately before each experiment. Control solutions contained the corresponding concentration of vehicle.

**Data and statistical analysis**

Electrophysiological and EEG data were analyzed using custom MATLAB scripts (MathWorks, USA) together with appropriate signal-processing routines. For *in vitro* experiments, tonic GABAA current density was calculated for individual VB TC neurons and expressed as pA/pF. The effect of MDL11,939 was assessed separately in NEC and GAERS neurons. The manuscript shows the individual neuronal measurements together with mean ± SEM.

For *in vivo* experiments, cumulative seizure time, mean seizure duration and seizure number were calculated for successive recording epochs and compared between vehicle- and MDL11,939-treated animals. Data are presented as mean ± SEM. Statistical significance was set at *P* < 0.05.
